# Sex-specific and pleiotropic effects underlying kidney function identified from GWAS meta-analysis

**DOI:** 10.1101/421552

**Authors:** Sarah E. Graham, Jonas B. Nielsen, Matthew Zawistowski, Wei Zhou, Lars G. Fritsche, Maiken E. Gabrielsen, Anne Heidi Skogholt, Ida Surakka, Damian Fermin, Sachin Kheterpal, Chad M. Brummett, Seunggeun Lee, Hyun Min Kang, Goncalo Abecasis, Solfrid Romundstad, Stein Hallan, Matthew G. Sampson, Kristian Hveem, Cristen J. Willer

## Abstract

Chronic Kidney Disease (CKD) is a growing health burden currently affecting 10-15% of adults worldwide. Estimated glomerular filtration rate (eGFR) as a marker of kidney function is commonly used to diagnose CKD. Previous genome-wide association study (GWAS) meta-analyses of CKD and eGFR or related phenotypes have identified a number of variants associated with kidney function, but these only explain a fraction of the variability in kidney phenotypes attributed to genetic components. To extend these studies, we analyzed data from the Nord-Trøndelag Health Study (HUNT), which is more densely imputed than previous studies, and performed a GWAS meta-analysis of eGFR with publicly available summary statistics, more than doubling the sample size of previous meta-analyses. We identified 147 loci (53 novel loci) associated with eGFR, including genes involved in transcriptional regulation, kidney development, cellular signaling, metabolism, and solute transport. Moreover, genes at these loci show enriched expression in urogenital tissues and highlight gene sets known to play a role in kidney function. In addition, sex-stratified analysis identified three regions (prioritized genes: *PPM1J, MCL1*, and *SLC47A1*) with more significant effects in women than men. Using genetic risk scores constructed from these eGFR meta-analysis results, we show that associated variants are generally predictive of CKD but improve detection only modestly compared with other known clinical risk factors. Collectively, these results yield additional insight into the genetic factors underlying kidney function and progression to CKD.

## Introduction

Chronic kidney disease (CKD) is a common condition affecting ∼11% of adults in Norway and ∼15% in the United States^1,2^. Due to specific comorbidities (namely diabetes) and an aging population, CKD is expected to continue to rise in global prevalence^3^. However, the prevalence varies with ethnicity and sex, with CKD being more common among females and African Americans^1^. Estimated glomerular filtration rate (eGFR) provides an assessment of kidney function and it is estimated based on serum creatinine levels with adjustment for age, race, and sex. eGFR levels below 60 mL/min/1.73m^2^characterize chronic kidney disease^4^, with varying severity classified by albuminuria and eGFR levels. A subset of individuals with CKD have accelerated renal function decline and progress to end stage renal disease (ESRD).

Several other diseases interact with kidney function. Chronic health conditions such as diabetes and hypertension directly influence the development of CKD, with environmental factors such as smoking accelerating disease progression^5^. Altered lipid levels, especially triglycerides, are also associated with CKD and contribute to the progression of cardiovascular disease^6^. Advanced stages of CKD/ESRD necessitate dialysis or transplantation and are associated with a greatly increased risk of cardiovascular disease and death^7^.

It has been estimated that about one third of the variation in eGFR levels can be attributed to genetic factors^8^, with the remaining variability due to environmental effects. Previous GWAS studies and meta-analyses have identified a number of loci associated with serum creatinine, eGFR, or CKD^9-18^. However, the introduction of denser imputation panels, including the Haplotype Reference Consortium^19^(HRC), and the recent rise in large-scale biobanks has enabled larger sample sizes and a greater number of variants than previously studied. Analysis of these new and more densely imputed datasets are expected to identify genetic regions influencing these traits not previously found. We analyzed samples from the Nord-Trøndelag Health Study (HUNT), imputed using a combined HRC and ancestry specific panel, for association with eGFR, creatinine, urea, and CKD, and performed meta-analysis of eGFR associations with three additional cohorts to uncover additional genetic variants contributing to kidney function.

## Results

### Meta-analysis of eGFR

Meta-analysis of up to 350,504 individuals from the HUNT Study, CKDGen Consortium, BioBank Japan, and the Michigan Genomics Initiative identified 147 loci associated with eGFR, of which 53 were novel (Table 1, **Supplementary Table 1, Supplementary Figure 1**). We prioritized genes belonging to several biological classes related to kidney function based on the consensus between the gene nearest to the index variant, identified significant missense variants that were either the lead variant or in moderate LD (r^2^> 0.3) with it, significantly colocalized eQTLs, and DEPICT gene prioritization results (**Supplementary Tables 2-4**). Prioritized genes at novel loci included genes involved in transcription (*CASZ1, NFE2L2, PPARGC1A, ZNF641, MED4, ZFHX3, ZGPAT, MAFF, MAMSTR*), cellular signaling and differentiation (*ACVR2B, DCDC2, GRB10, NRG1, THADA, TRIB1, PTPN3, FAM53B*), metabolism (*L2HGDH, XYLB*), solute carrier genes (*SLC25A43, TPCN2, KCNMA1, MFSD6*), and a gene related to AB antigen blood types (*ABO*). As no similarly-sized cohort with eGFR measurements was available for direct replication, we instead tested for association of the index variants in kidney related traits in the UK Biobank (CKD, hypertensive CKD, renal failure, acute renal failure, renal failure NOS, renal dialysis, or other disorders of kidney and ureters). Thirteen of the 48 novel variants and 36 of the 85 lead variants in known loci that were available in the UK Biobank were at least nominally associated with one or more UK Biobank kidney-related phenotypes (**Supplementary Table 5**). In addition, we compared the results from the current meta-analysis with previously reported eGFR index variants^9-12,14,15,17^. Of the 127 previously reported index variants for eGFR, 125 were at least nominally significant in the present meta-analysis (**Supplementary Table 6**) and 63% reached genome-wide significance (p-value < 5×10^−8^). Excluding the previously published datasets, 56 of the 118 available variants were at least nominally significant in meta-analysis of HUNT and MGI alone (**Supplementary Table 6**).

**Table 1:** Lead Variants for Novel eGFR Loci from Meta-analysis

| Chr | Pos (hg19) | rsID | Ref | Alt | Freq<sup>a</sup> | P-value | Direction<sup>a</sup> | Prioritized Gene |
| --- | --- | --- | --- | --- | --- | --- | --- | --- |
| 1 | 10733081 | rs284316 | T | C | 0.3331 | 1.50 x 10<sup>-9</sup> | + | *CASZ1* |
| 1 | 100808363 | rs11166440 | A | G | 0.4119 | 7.07 x 10<sup>-9</sup> | - | *CDC14A* |
| 1 | 180905694 | rs3795503 | T | C | 0.6017 | 7.54 x 10<sup>-13</sup> | - | *KIAA1614* |
| 1 | 227085824 | rs1800674 | A | G | 0.5615 | 4.23 x 10<sup>-8</sup> | - | *PSEN2* |
| 2 | 18679586 | rs10856778 | C | G | 0.8311 | 1.04 x 10<sup>-9</sup> | + | *LOC105373454* |
| 2 | 43441169 | rs35136921 | T | C | 0.4564 | 2.74 x 10<sup>-11</sup> | + | *THADA* |
| 2 | 54574942 | rs1405833 | C | G | 0.2733 | 5.10 x 10<sup>-10</sup> | - | *C2orf73* |
| 2 | 178146362 | rs17581525 | C | G | 0.1889 | 6.11 x 10<sup>-11</sup> | + | *NFE2L2* |
| 2 | 191278341 | rs6725814 | A | G | 0.261 | 3.39 x 10<sup>-8</sup> | + | *MFSD6* |
| 2 | 230612451 | rs6756038 | A | G | 0.7353 | 4.72 x 10<sup>-9</sup> | - | *TRIP12* |
| 3 | 38479475 | rs7429308 | T | C | 0.4951 | 4.90 x 10<sup>-11</sup> | - | *ACVR2B, XYLB* |
| 3 | 193816778 | rs10933714 | A | T | 0.5232 | 2.81 x 10<sup>-9</sup> | - | *LOC102724877* |
| 3 | 195477791 | rs2291652 | A | G | 0.4385 | 2.36 x 10<sup>-8</sup> | - | *MUC4* |
| 4 | 23758662 | rs73243607 | T | C | 0.962 | 3.37 x 10<sup>-8</sup> | - | *PPARGC1A* |
| 5 | 107459529 | rs12652687 | T | C | 0.1447 | 3.61 x 10<sup>-8</sup> | - | *FBXL17* |
| 6 | 24354045 | rs3765502 | T | C | 0.2351 | 4.01 x 10<sup>-8</sup> | - | *DCDC2* |
| 6 | 107172979 | rs7766720 | T | C | 0.1424 | 4.13 x 10<sup>-8</sup> | - | *LINC02532* |
| 7 | 50737852 | rs73116822 | T | C | 0.9282 | 2.71 x 10<sup>-9</sup> | + | *GRB10* |
| 7 | 56072841 | rs4948100 | T | C | 0.3796 | 1.17 x 10<sup>-8</sup> | + | *GBAS* |
| 7 | 128737958 | rs56088330 | A | T | 0.6814 | 8.12 x 10<sup>-10</sup> | + | *LOC112267982* |
| 8 | 9074223 | rs7006504 | T | C | 0.297 | 1.91 x 10<sup>-9</sup> | - | *RP11-10A14.5* |
| 8 | 32399662 | rs4489283 | T | C | 0.6897 | 2.47 x 10<sup>-9</sup> | + | *NRG1* |
| 8 | 126477978 | rs2001945 | C | G | 0.4361 | 4.37 x 10<sup>-11</sup> | + | *TRIB1* |
| 8 | 134332960 | rs10283362 | T | C | 0.8607 | 2.04 x 10<sup>-8</sup> | - | *RPL32P20* |
| 9 | 34130435 | rs61237993 | A | G | 0.654 | 1.85 x 10<sup>-8</sup> | - | *DCAF12* |
| 9 | 112206404 | rs10816812 | A | T | 0.6295 | 1.00 x 10<sup>-8</sup> | + | *PTPN3* |
| 9 | 136146597 | rs550057 | T | C | 0.7393 | 1.58 x 10<sup>-9</sup> | - | *ABO* |
| 9 | 139107879 | rs11103387 | T | C | 0.7296 | 8.07 x 10<sup>-10</sup> | - | *QSOX2* |
| 10 | 35171118 | rs11010013 | A | G | 0.6928 | 1.50 x 10<sup>-8</sup> | - | *SS18L2P1* |
| 10 | 79253261 | rs3127447 | A | C | 0.3283 | 3.04 x 10<sup>-8</sup> | + | *KCNMA1* |
| 10 | 94810665 | rs856534 | A | G | 0.4342 | 1.23 x 10<sup>-9</sup> | + | *EXOC6* |
| 10 | 126418782 | rs11245344 | T | C | 0.4673 | 3.37 x 10<sup>-11</sup> | + | *FAM53B* |
| 11 | 68883556 | rs7131509 | T | C | 0.5311 | 2.76 x 10<sup>-10</sup> | - | *TPCN2, LOC107984345* |
| 12 | 48736985 | rs2732481 | T | G | 0.2444 | 1.82 x 10<sup>-9</sup> | - | *ZNF641* |
| 13 | 48654455 | rs9534949 | C | G | 0.6165 | 4.84 x 10<sup>-9</sup> | + | *MED4* |
| 14 | 50735947 | rs72683923 | T | C | 0.0109 | 2.98 x 10<sup>-8</sup> | + | *L2HGDH* |
| 15 | 39274261 | rs8026431 | A | C | 0.6364 | 3.13 x 10<sup>-9</sup> | - | *LOC105370781* |
| 15 | 57830151 | rs117047297 | T | C | 0.9836 | 3.16 x 10<sup>-8</sup> | - | *CGNL1* |
| 15 | 63634405 | rs1075456 | T | G | 0.6601 | 3.05 x 10<sup>-9</sup> | - | *CA12* |
| 15 | 67561355 | rs12443279 | C | G | 0.3495 | $3.03 \times 10^{-8}$ | - | *IQCH* |
| 16 | 69718112 | rs77944668 | A | G | 0.6984 | $1.45 \times 10^{-8}$ | - | *NFAT5* |
| 16 | 73024276 | rs1858800 | T | C | 0.761 | $1.82 \times 10^{-8}$ | - | *ZFXH3* |
| 16 | 79938996 | rs35286975 | C | G | 0.7933 | $1.94 \times 10^{-10}$ | - | *LINC01229* |
| 17 | 34950239 | rs12937411 | T | C | 0.6134 | $1.62 \times 10^{-10}$ | - | *MYO19, DHRS11* |
| 19 | 18384950 | rs1075403 | T | G | 0.6992 | $1.57 \times 10^{-10}$ | - | *KIAA1683* |
| 19 | 49217305 | rs281386 | A | G | 0.591 | $4.77 \times 10^{-8}$ | + | *MAMSTR* |
| 20 | 62336334 | rs1758206 | T | C | 0.8245 | $1.51 \times 10^{-9}$ | + | *ZGPAT, LIME1* |
| 21 | 16582710 | rs56038390 | A | G | 0.3014 | $1.28 \times 10^{-9}$ | - | *LOC105369292* |
| 21 | 35356706 | rs2834317 | A | G | 0.8977 | $1.58 \times 10^{-8}$ | + | *LOC105372790* |
| 22 | 38600542 | rs2267373 | T | C | 0.3949 | $2.65 \times 10^{-10}$ | + | *MAFF* |
| X | 18597869 | rs4825261 | A | C | 0.7148 | $1.36 \times 10^{-9}$ | - | *CDKL5* |
| X | 118630622 | rs454741 | A | G | 0.641 | $5.15 \times 10^{-10}$ | - | *LOC107985719, SLC25A43, CXorf56* |
| X | 133808916 | rs11796053 | A | G | 0.4934 | $4.93 \times 10^{-8}$ | + | *PLAC1* |

<sup>a</sup>Reported frequency and direction of effect is with respect to the alternative allele

**Table 2:**
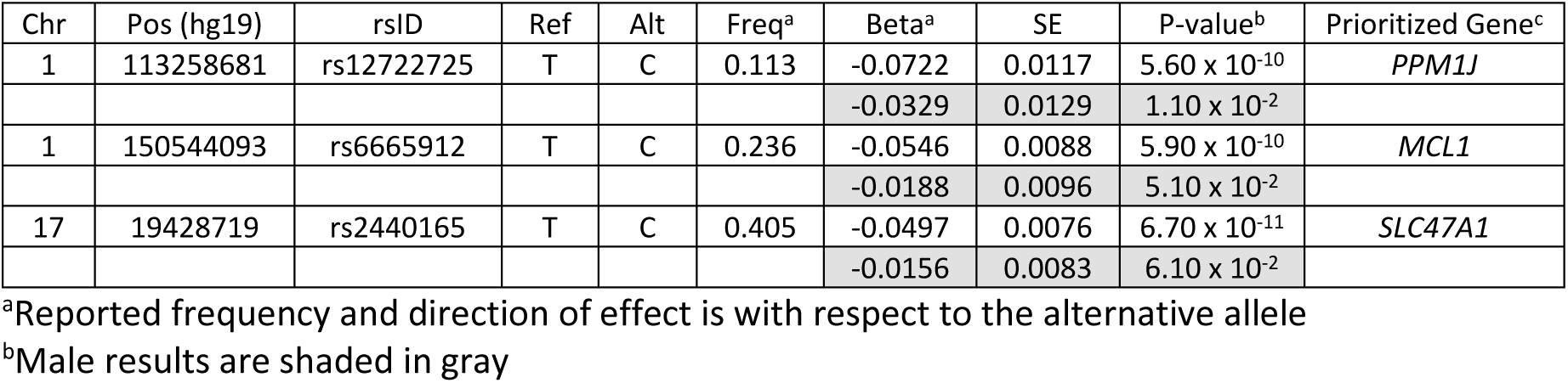
Lead Variants for Loci Showing Differences between Males and Females in HUNT Study

| Chr | Pos (hg19) | rsID | Ref | Alt | Freq<sup>a</sup> | Beta<sup>a</sup> | SE | P-value<sup>b</sup> | Prioritized Gene<sup>c</sup> |
| --- | --- | --- | --- | --- | --- | --- | --- | --- | --- |
| 1 | 113258681 | rs12722725 | T | C | 0.113 | -0.0722 | 0.0117 | 5.60 x 10<sup>-10</sup> | *PPM1J* |
|  |  |  |  |  |  | -0.0329 | 0.0129 | 1.10 x 10<sup>-2</sup> |  |
| 1 | 150544093 | rs6665912 | T | C | 0.236 | -0.0546 | 0.0088 | 5.90 x 10<sup>-10</sup> | *MCL1* |
|  |  |  |  |  |  | -0.0188 | 0.0096 | 5.10 x 10<sup>-2</sup> |  |
| 17 | 19428719 | rs2440165 | T | C | 0.405 | -0.0497 | 0.0076 | 6.70 x 10<sup>-11</sup> | *SLC47A1* |
|  |  |  |  |  |  | -0.0156 | 0.0083 | 6.10 x 10<sup>-2</sup> |  |

<sup>a</sup>Reported frequency and direction of effect is with respect to the alternative allele<sup>b</sup>Male results are shaded in gray

**Figure 1:**
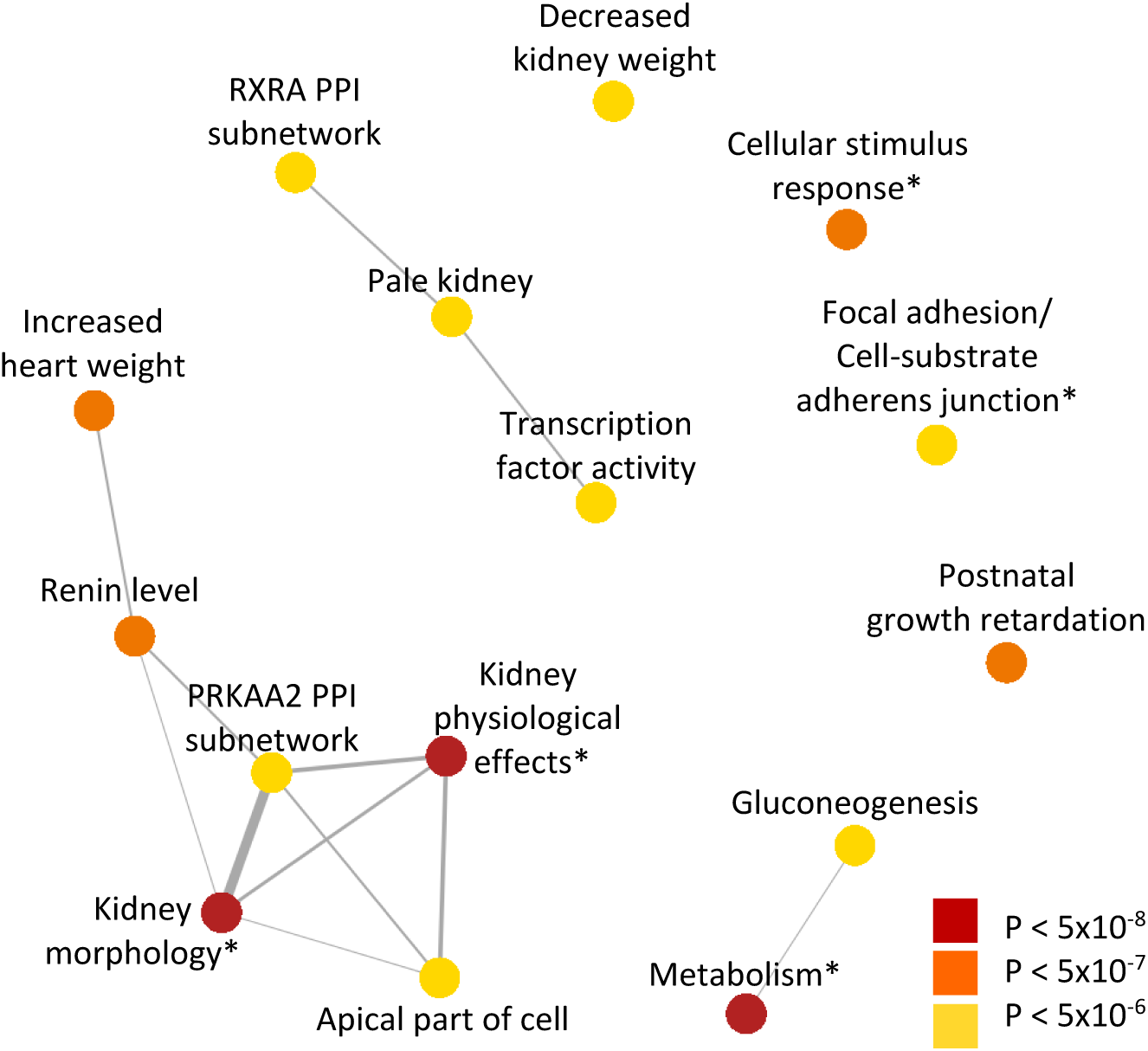
Gene sets prioritized from eGFR meta-analysis. DEPICT analysis of eGFR meta-analysis results identifies significant gene sets associated with kidney function and metabolic processes. Overlap between gene sets is depicted by the width of connecting lines. *Denotes collapsed gene sets.

### Kidney-Specific eQTL Associations

To identify variants that may be acting through regulation of gene expression in a kidney-specific manner, we selected eGFR index variants which were significant eQTLs (p-value < 6.7×10^−6^, Bonferroni correction for 51 tissue types and 147 index variants) for a given gene in kidney cortex^20^, glomerulus^21^, or tubulointerstitium, but not in other tissues in GTEx^22^. This analysis identified five genes whose expression was associated with the eGFR index variants only in kidney tissues: *APOD, CDKL5, DPEP1, FGF5*, and *TFDP2* (**Supplementary Table 7**). Of these, *FGF5* and *CDKL5* kidney-eQTLS showed significant colocalization with the eGFR association.

### DEPICT Analysis

DEPICT analysis was performed to identify tissues and gene sets enriched for genes in the loci identified from eGFR meta-analysis. Consistent with the role of the identified genes in kidney function, the only significant tissues (p-value < 2.39×10^−4^) identified by DEPICT were the urinary tract (p-value = 2.1×10^−6^) and kidney urogenital system (p-value = 2.8×10^−6^) (**Supplementary Table 8**). Significant gene sets identified by DEPICT primarily included those associated with kidney morphology, the activity of transport channels, and with monosaccharide metabolic processes as shown in **Figure 1** (**Supplementary Figure 2**, **Supplementary Table 9**).

### Overlap with Related Traits

As individuals with CKD often have coexisting heart disease or diabetes, we examined the identified eGFR variants for evidence of pleiotropic effects. Lookup of the index variants across 26 cardiovascular and diabetes-related phenotypes in UK Biobank, excluding individuals with CKD, identified 7 phenotypes for which a subset of the index variants was also genome-wide significant (diabetes, coronary atherosclerosis, hypertension, essential hypertension, other disorders of circulatory system, pulmonary heart disease, and phlebitis and thrombophlebitis), and several additional phenotypes for which the eGFR variants showed nominal significance (**Supplementary Tables 5**,**10**). Colocalization analysis with these phenotypes identified 7 loci (prioritized genes: *FGF5, PRKAG2, TRIB1, LOC101928316, L2HGDH, UMOD/PDILT, LOC105371257*) having significantly colocalized association signals with hypertension, essential hypertension, and/or coronary atherosclerosis and 1 locus (prioritized gene: *GCKR*) that colocalized with association of type 2 diabetes (**Supplementary Table 5**). Six of the seven index variants within loci that showed significant colocalization with the cardiovascular traits were associated with essential hypertension and/or hypertension, underscoring the connection between high blood pressure and CKD. In addition, the index variants were examined for association with 1,403 traits phenome-wide (without exclusion of CKD cases). As shown in **Figure 2**, the index variants are significantly associated (p-value < 5×10^−8^) with additional traits including hypothyroidism and disorders of lipid metabolism.

**Figure 2:**
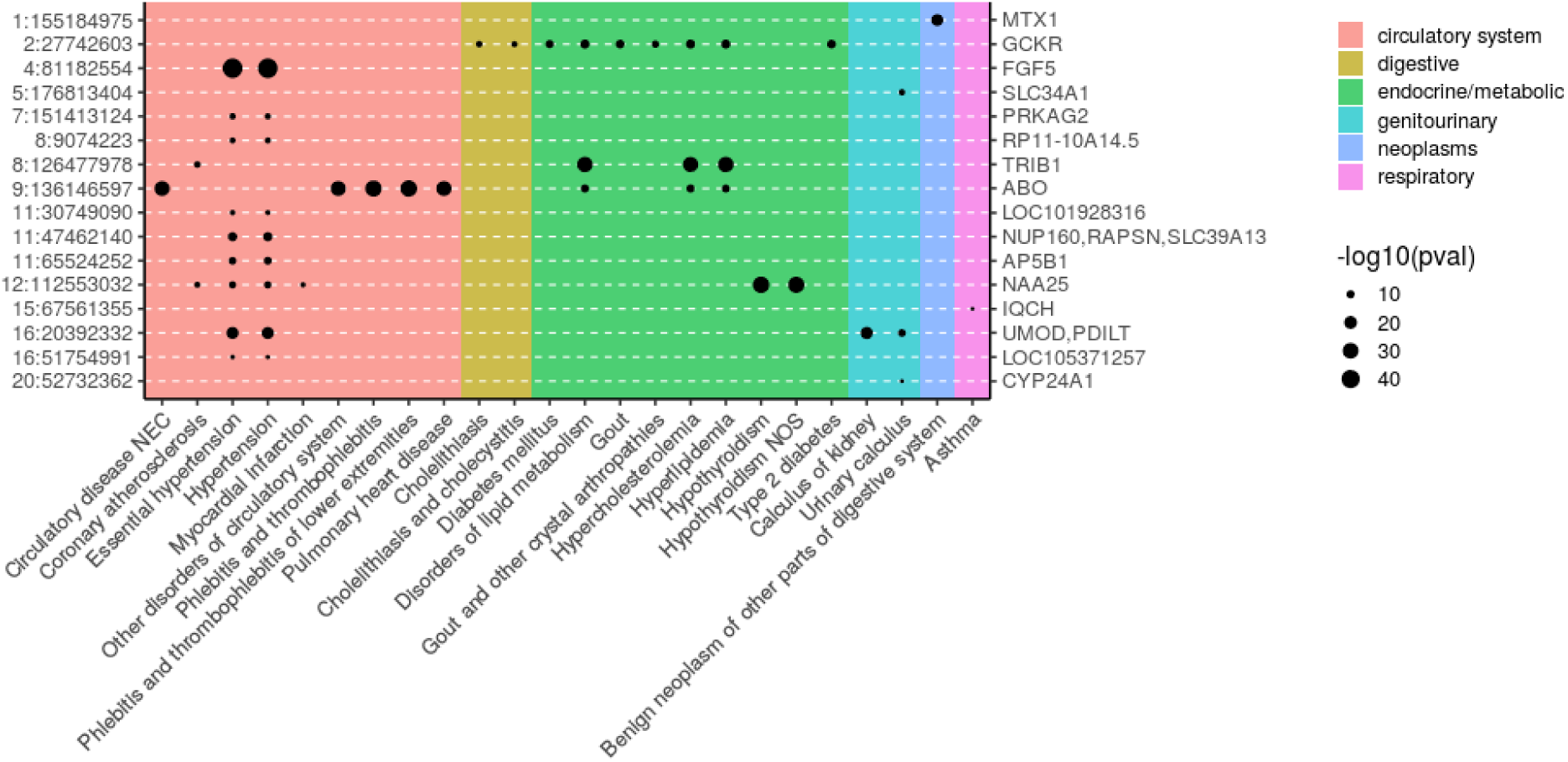
Pleiotropic associations of eGFR Index Variants. Index variants, given as chromosome:position on the left axis and prioritized gene on the right, from eGFR meta-analysis showing significant associations (p-value < 5×10^−8^) with at least one additional phenotype in UK Biobank (N^max^ = 408,961). 127 variants were tested for association with 1,403 phenotypes.

### Construction of genetic risk scores

We developed genetic risk scores (GRS) from the meta-analysis results to assess the relationship between the identified variants and the likelihood of having CKD. Several p-value and r^2^clumping thresholds were tested to select variants for inclusion in the GRS, but all yielded relatively similar predictions of CKD status (AUC range: 0.500-0.533). The best prediction was obtained using all independent markers (r^2^< 0.2) with p-value < 5×10^−6^and the European and East Asian subsets of 1000 Genomes for LD clumping. These scores were then tested as predictors of CKD in UK Biobank. The GRS alone was associated with CKD (p-value = 2.13×10^−11^), but did not improve prediction of CKD status compared to birth year and sex alone (AUC: 0.694), or birth year, sex, and CKD clinical risk factors (diabetes, hypertension, and hyperlipidemia) (AUC: 0.864). Inclusion of the GRS in addition to birth year, sex, and clinical risk factors provided the best predictor of CKD of the tested models (AUC: 0.865). We also tested prediction of CKD using the best-performing GRS from the overall meta-analysis (without birth year or additional risk factors) separately in men and women. The GRS was slightly more predictive in women (AUC: 0.543) than in men (AUC: 0.527), possibly due to differing lifestyle or hormonal factors between the sexes influencing development of CKD. These results show that the variants identified from association studies of eGFR are correlated with the presence of CKD on a population level, but are not sufficient to identify individuals with CKD from those without. This is consistent with findings from prior studies examining GRS of eGFR^23^.

### Sex-specific Analysis in HUNT

We aimed to determine whether any variants showed sex-specific association as there are known differences in the prevalence of CKD between men and women. Association tests of eGFR in HUNT stratified by sex identified three loci (Figure 3, **Supplementary Figure 3**) that had significantly different effect sizes between men and women (p-value for difference < 0.017, **Table 2**). At these loci, we prioritized 3 candidate functional genes, *PPM1J, MCL1*, and *SLC47A1*, based on significant colocalization of eQTL and eGFR association signals and high linkage disequilibrium with missense variants. Interestingly, lookup of these variants in UK Biobank for association with other phenotypes (http://pheweb.sph.umich.edu:5003) identified significant associations with rs6665912 (*MCL1*) for increased occurrence of chest pain (p-value = 4.6×10^−6^) and coronary atherosclerosis (p-value = 2.3×10^−5^). Sex-stratified analysis of chest pain and coronary atherosclerosis after exclusion of CKD cases in UK Biobank showed a consistent sex difference for both traits for this variant, though with opposite sex-specificity to that seen for eGFR (chest pain: p-value_men_ = 6.7×10^−5^, p-value_women_ = 0.041, coronary atherosclerosis: p-value_men_ = 1.9×10^−4^, p-value_women_ = 0.021).

**Figure 3:**
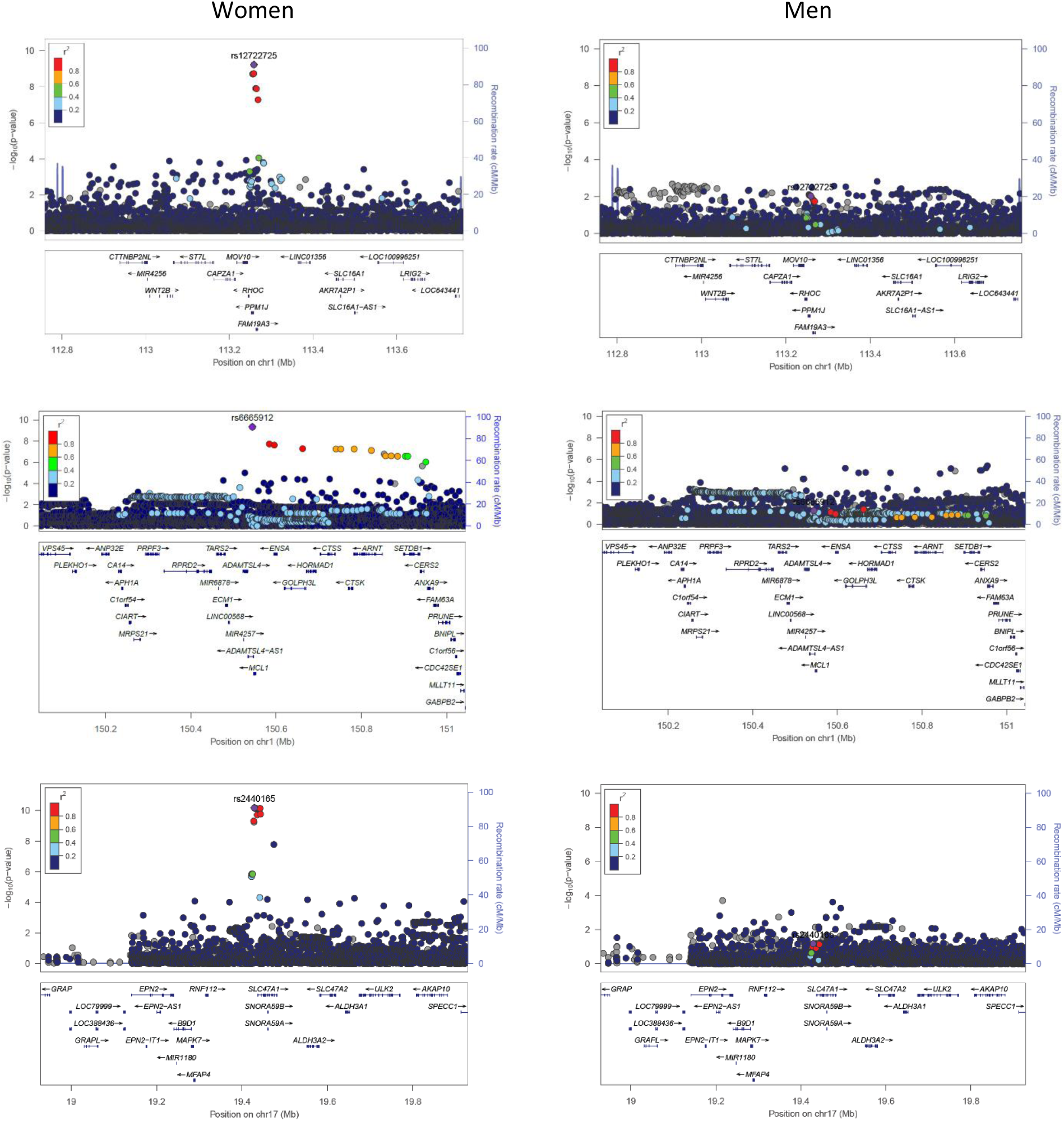
Locus zoom plots of regions showing differential association between men and women. eGFR meta-analysis results in HUNT stratified by sex were filtered to identify regions significant in one sex but near-nominal significance in the other.

We obtained the summary statistics from the CKDGen consortium sex-stratified eGFR analysis in order to test these variants for replication^14^. As the CKDGen consortium results were imputed using HapMap-II, we performed meta-analysis of the HUNT and CKDGen sex-specific results and then assessed the identified lead variants or available LD proxies for consistent association. After meta-analysis, both rs6665912 (*MCL1*) and the proxy variant of rs2440165 (rs2453580; r^2^= 0.997, *SLC47A1*) showed greater significance in women than in men (rs6665912: p-value_women_= 3.67×10^−7^, p-value_men_= 0.059, rs2453580: p-value_women_ = 3.36×10^−13^, p-value_men_ = 0.0030). However, rs6665912 was not significant in the CKDGen data alone. We were unable to test rs12722725 (*PPM1J*) for replication, as neither this variant nor any LD proxies with r^2^> 0.6 were available in the HapMap-II imputed results.

### Analysis in HUNT

Within the HUNT dataset alone, a total of 28 significant eGFR loci were identified, of which 1 was novel (**Supplementary Tables 11,12, Supplementary Figure 4**). The novel variant, prioritized gene *CDKL5*, was also identified through the combined meta-analysis. Step-wise conditional analysis was performed and 3 additional variants within the identified eGFR loci reached genome-wide significance (**Supplemental Table 13**). Analysis of urea and creatinine identified 4 and 25 previously known loci, respectively, while analysis of CKD did not identify any significant variants with MAF > 0.5%. In addition, analysis of custom variants included in the genotyping array for a subset of individuals (N= 37,472) identified one rare stop-gain variant in the known locus *PKD2* associated with creatinine (**Supplementary Table 12**).

## Discussion

In summary, analysis of the HUNT biobank and meta-analysis of eGFR across more than 350,000 individuals identified 147 loci, of which 53 were novel. The identified loci give new insight into the genes underlying kidney function and development of CKD. In support of this, many of the prioritized genes cluster into known kidney associated pathways. For example, Wnt signaling has been implicated in kidney development and disease^24^. FAM53B controls β-catenin nuclear localization^25^, a key component of Wnt signaling. β-catenin translocation into the nucleus allows for activation of target genes via interaction with TCF transcription factors^26,27^. Variants in *FAM53B* were associated with changes in eGFR, which is potentially due to altered Wnt signaling and expression of target genes. Similarly, *DCDC2* variants were also associated with eGFR levels. Knockdown or overexpression of DCDC2 is known to alter β-catenin activation of TCF transcription factors^28^, thereby altering Wnt signaling. Likewise, variants in genes associated with the epidermal growth factor receptor (ErbB) family were also observed. ErbB receptors are involved in kidney development^29^, control of solute levels (eg. Ca, Na)^30,31^, and play a role in hypertension^32,33^. Our eGFR association results identified variants in two genes (*NRG1* and *MUC4*) known to bind to ErbB receptors. NRG1 binds to ErbB3 and ErbB4 while the beta chain of Mucin-4 (*MUC4*) interacts with ErbB2^34^. Lastly, we identified variants associated with both decreased eGFR and increased tubulointerstitial kidney CDKL5 expression. CDKL5 overexpression has been shown to impair ciliogenesis^35^. Defects in cilia are known to cause polycystic kidney disease and nephronophthisis, among other disorders^36^. These clues provide an initial link to how these identified genetic regions may lead to changes in kidney function.

Experimental evidence also supports hormonal regulation of *MCL1* and *SLC47A1* expression, two of the genes prioritized from the sex-stratified analysis of eGFR. For example, Mcl-1 (encoded by *MCL1*) functions as a regulator of apoptosis and has been extensively studied due to its role in carcinogenesis. In estrogen-receptor positive breast cancer cell lines, estrogen increased expression of Mcl-1^37^. *SLC47A1* is also known as *MATE1* (multidrug toxin and extrusion protein 1). Experimental studies of MATE1 identified higher levels of expression in the kidneys of 30-45 day old male mice compared to female^38^. Furthermore, He at al. found that kidney expression of MATE1 could be modified by treatment with testosterone or estradiol, compared to olive oil as a control^39^.

It is also interesting to consider the interplay between kidney function and other related traits based on the overlap between identified genetic regions. For example, the *ABO* gene was prioritized based on eGFR meta-analysis results. *ABO* encodes a protein that is responsible for determination of an individual’s ABO blood type. Associations near this gene have been previously identified for other phenotypes, including LDL and total cholesterol^40^, coronary artery disease^41^, and type 2 diabetes^42^. As diabetes is a significant risk factor for the development of CKD, these shared associations may help to identify potential common mechanisms. Comparison of association results with cardiovascular related traits also identified shared associations with hypertension, the second major risk factor for CKD. Six of the 147 loci identified from meta-analysis showed significant colocalization with hypertension, which may help to identify additional shared pathways between high blood pressure and kidney function.

While additional studies are needed to understand eGFR associations that are specific to disease subtypes, the present results build upon previous studies to increase the number of eGFR associated loci and highlight pleiotropic associations with cardiovascular disease. Follow-up experimental studies are needed to validate the role of the identified genes in kidney function, and additional genetic studies are needed to verify these associations in more diverse cohorts. Nevertheless, these results identify additional genes that are likely involved in regulating kidney function and may help to identify new therapeutic targets or diagnostic measures for progression to chronic kidney disease.

## Methods

### Description of Cohorts

#### HUNT

The HUNT study^43^is a longitudinal, repetitive population-based health survey conducted in the county of Nord-Trøndelag, Norway in which kidney-related phenotypes have not previously been tested for genetic association. Since 1984, the entire adult population in the county has been examined three times, through HUNT1 (1984-86), HUNT2 (1995-97), and HUNT3 (2006-08). A fourth survey, HUNT4 (2017-2019), is ongoing. Approximately 120,000 individuals have participated in HUNT1-HUNT3 with extensive phenotypic measurements and biological samples. A subset of these participants have been genotyped using Illumina HumanCoreExome v1.0 and 1.1 and imputed with Minimac3 using a combined HRC and HUNT-specific WGS reference panel. Variants with imputation r^2^< 0.3 were excluded from further analysis. We analyzed available kidney related phenotypes within the HUNT study, including creatinine (N = 69,591), eGFR (N = 69,591), urea (N = 20,700), and CKD (N_cases_ = 2044 and N_controls_= 65,575). eGFR values were calculated using the MDRD equation^44,45^. CKD status was taken from ICD-9 codes 585 and 586 and ICD-10 code N18. Association testing of quantitative traits was performed using BOLT-LMM^46^v2.2 on the inverse-normalized residuals of the traits adjusted for genotyping batch, sex, 4 principle components, and age. Association testing for CKD was performed using SAIGE^47^with sex, 4 principle components, and birth year as covariates. Associations stratified by sex for eGFR were also performed. For the stratified analyses, phenotypes were separately inverse-normalized and were adjusted for batch, age, and 4 principle components. Linkage disequilibrium within HUNT was calculated using PLINK v1.90^48^. Conditional analysis for eGFR was performed within the HUNT dataset using BOLT-LMM v.2.3.1, conditioning on the lead variant within the identified loci until no variants with MAF > 0.5% had p-value < 5×10^−8^. To identify sex-specific effects, loci were identified separately in men and women and were filtered to those that were significant in one sex but near nominal significance in the other. Differences in effect sizes between males and females were then tested using Z = (β_M_ -β_W_)/(SE_M_ ^2^ + SE _W_ ^2^ – 2rSE_M_ SE_W_)^0.5^, where r is the Pearson correlation for male and female effect sizes across all variants^49^. Significance between sexes was determined using Bonferroni correction for the number of tested loci.

#### BBJ

BioBank Japan (BBJ) is a registry of patients from 12 medical centers across Japan who are diagnosed with at least one of 47 common diseases^50^. Summary statistics for 58 quantitative traits, including eGFR, are publicly available^17^. Participating individuals were genotyped with either the Illumina HumanOmniExpressExome BeadChip or HumanOmniExpress and HumanExome BeadChips. Imputation was performed with Minimac using the East Asian reference panel from 1000 Genomes phase 1^51^. Variants with imputation r^2^< 0.7 were excluded. BBJ eGFR values were calculated using the Japanese ancestry modified version of the CKD-EPI equation^52^and were available on 143,658 of those enrolled. Individuals with eGFR values of less than 15 mL/min/1.73 m^2^were excluded from analysis. Values were standardized using rank-based inverse normalization. Association analysis was performed using mach2qtl with sex, age, the top 10 principle components, and disease status of all studied diseases (N = 47) included as covariates.

### CKDGen Consortium

The CKDGen consortium includes meta-analysis results from 33 individual studies of European ancestry (N = 110,527) that were imputed with the 1000 Genomes phase I reference panel^9^. Summary statistics for eGFR were taken from the published dataset (ckdgen.imbi.uni-freiburg.de). Detailed descriptions of individual cohorts are available^9^. Briefly, each group generated association statistics based on the natural log of eGFR using age and sex as covariates. eGFR was estimated from creatinine levels using the MDRD equation^44,45^. Variants with imputation quality ≤ 0.4, and those found in less than half of individuals were excluded from further analysis. Meta-analysis was performed using the inverse-variance method in METAL^53^. Pre and post meta-analysis genomic control (GC) correction was performed.

#### MGI

The Michigan Genomics Initiative (MGI) is a repository of patient electronic health and genetic data at Michigan Medicine^54^(N = 26,738). MGI participants are recruited primarily through pre-surgical encounters at Michigan Medicine and consent to linking of genetic and clinical data for research purposes. DNA was extracted from blood samples and then participants were genotyped using Illumina Infinium CoreExome-24 bead arrays. Genotype data was then imputed to the Haplotype Reference Consortium using the Michigan Imputation Server, providing 17 million imputed variants after standard quality control and filtering. eGFR values were computed using the CKD-EPI equation from creatinine values. The mean eGFR value was used for individuals having more than one eGFR measurement. Mean eGFR was then regressed on sex, current age, array version and PC1-4 and the subsequent residuals were inverse-normalized. Single-variant association testing of the inverse normalized residuals was performed in *epacts* using a linear regression model.

#### Meta-analysis

Meta-analysis was performed using the p-value based approach in METAL^53^. This approach was chosen to account for differing units between the effect sizes of the CKDGen (log-transformed) and MGI/BBJ/HUNT (inverse-normalized) summary statistics. This approach was validated by comparison to traditional standard error-based meta-analysis of the cohorts with available inverse-normalized summary statistics, and showed extremely high correlation of p-values (Pearson r= 0.966, **Supplementary Figure 5**). Summary statistics from contributing studies were GC corrected prior to meta-analysis. Lead index variants were determined as the most significant variant in ± 1 Mb windows that were found in at least 2 studies. Adjacent windows were merged if the LD r^2^between lead variants was ≥ 0.2. Identified variants were considered to be novel if the most significant variant was more than 1 Mb away from previously reported lead variants. Linkage disequilibrium between variants was calculated using LDlink^55^ or PLINK^48^with the European and East Asian 1000 Genomes Phase III reference panels^56^.

#### Variant and Gene Annotation

Variants were annotated using WGSA^57^and dbSNP^58^. Annotation of variants with associated biological processes was performed using the UniProt^59^and NCBI gene (https://www.ncbi.nlm.nih.gov/gene) databases. Genes for identified loci were prioritized based on the consensus between the nearest gene, significantly colocalized eQTLs, missense variants within 1 Mb and in moderate LD (r^2^> 0.3) with the lead variant, and the gene prioritized by Data-driven Expression-Prioritized Integration for Complex Traits (DEPICT)^60^(**Supplementary Figure 6**). In cases where there was not consensus between the different annotation methods, the gene was prioritized as the nearest gene. DEPICT analysis was performed using the DEPICT 1.1 1000 Genomes version. Variants from meta-analysis that were found in two or more studies with p-value < 5×10^−8^were included. LD information from the European and East Asian subsets of 1000 Genomes was used to construct loci within DEPICT. Significance of DEPICT results was determined using Bonferroni correction across the number of tissues or gene-sets tested. Gene sets with more than 25% overlap were collapsed into a single set for construction of the network diagram, as previously done^61^. Colocalization analysis of the kidney association results with GTEx V7^22^, NephQTL^21^, and Ko et al.^20^eQTL data was performed using the R package coloc^62^. Priors for p1, p2, and p12 within the coloc analysis were set to 1×10^−4^, 1×10^−4^, and 1×10^−6^, respectively. Following the criteria published by Giambartolomei et al.^62^, eQTLs were considered to colocalize with the kidney association results if the posterior probability (PP) for a shared variant was > 80%.

### Comparison with Related Traits

Association results for phenome-wide lookup were taken from analysis of UK Biobank^63^using SAIGE^47^, which accounts for relatedness and population stratification by using a relationship matrix. CKD cases were excluded from the analysis of cardiovascular and diabetic traits as has been previously suggested for identifying pleiotropic effects^54^. Colocalization analysis was performed using the R package coloc, with the priors for p1, p2, and p12 set to 1×10^−4^, 1×10^−4^, and 1×10^−6^, respectively. Genetic regions for colocalization testing were defined as the most significant variant in each locus ± 500 kb. Variants were considered to colocalize if the probability for a common variant was greater than 80%.

### Genetic Risk Scores

Variants were selected for inclusion in the genetic risk score (GRS) using the clumping procedure in PLINK^48^with varying r^2^thresholds of 0.2, 0.4, 0.6, and 0.8 and p-value thresholds of 5×10^−8^, 5×10^−6^, 5×10^−4^, and 5×10^−3^on the meta-analysis results from all cohorts. As the meta-analysis included both European and East Asian samples but the validation set included primarily European samples, we separately constructed the GRS using either the European only or European and East Asian subsets of 1000 Genomes Phase 3 for clumping in PLINK. Effect sizes were estimated from meta-analysis of the BioBank Japan, MGI, and HUNT results only due to the differing effect size units of the CKDGen consortium results. GRS were then calculated within UK Biobank as the sum of risk alleles carried by each individual weighted by the effect size of each variant. As decreased eGFR is predictive of increased CKD, the negative value of the resulting risk score was used for further analysis. GRS were then tested as predictors of CKD, either alone or as a logistic model including birth year, sex, and GRS or birth year, sex, GRS, and diabetes, hypertension, and hyperlipidemia status. When fitting the logistic model for prediction of CKD, individuals in UK Biobank were randomly split into two halves, with one half of individuals used for model fitting and the other half used for testing of the model. Prediction ability was assessed by area under the ROC curve (AUC).

## Acknowledgments

We thank all research participants in the HUNT study and in all included studies for their dedication towards improving human health. We also thank Whitney Hornsby and Bethany Klunder for project management. We thank the CKDGen Consortium for publicly sharing CKD summary statistics and for making available sex-specific results for a subset of variants following our request. Funding for this study was provided by the National Institutes of Health to CJW (R01 HL127564, R35 HL135824). MGS is supported by the Charles Woodson Clinical Research Fund, the Ravitz Foundation, and by National Institutes of Health RO1-DK108805. This research has been conducted using the UK Biobank Resource under application number 24460.

